# Inflammasome Activation in Cutaneous Squamous Cell Carcinoma

**DOI:** 10.1101/2025.04.29.650608

**Authors:** Kush R Patel, Raphael R Shu, Ruochong Wang, Khushi Tekale, Michael Hamersky, Adelaida B Perez, David Solano, Nooreen Syed, Connor E Stewart, Joseph Lee, Allison M Hanlon, Travis W Blalock, Chaoran Li, Lindsey Seldin

## Abstract

Epithelia maintain their barrier function by exploiting the proliferative and plastic properties of stem cells to promote continual tissue regeneration. Consequently, however, these features make epithelia prone to tumorigenesis. Skin is the largest epithelial barrier and the source of cutaneous cancers, the deadliest of which are cutaneous squamous cell carcinomas (cSCCs). Despite the pervasiveness of these cancers, however, the molecular mechanisms employed by stem cells and their microenvironment to promote skin tumor development remain poorly defined. Our previous work revealed that genotoxic damage to normal skin activates inflammasome signaling crosstalk between epidermal epithelia and fibroblasts to promote epithelial stem cell hyperproliferation and fate misspecification. We hypothesized that these phenomena would also be featured during skin cancer development. Using mouse and human skin disease specimens, we determined that in vivo stem cell misspecification is a generalizable feature across diverse pathophysiological skin conditions, including cutaneous cancers, but is not present in normal high proliferative contexts. Strikingly, in vivo inflammasome activation was observed in both the epithelial and dermal compartments of cSCC tumors but was not evident in other skin pathologies. Probing further into the mechanism, we revealed that inflammasome pathway activation in cSCC is epithelial autonomous but non-cell autonomous. Furthermore, fibroblasts juxtaposed to the cSCC epithelial tumor interface exhibited population expansion as well as IL-1 signaling activation, which was absent in overlying epithelia. Based on these findings, we propose a model whereby epithelial-to-fibroblast inflammasome crosstalk initiates a fibroblast feed-forward IL-1 signaling loop that augments the tumor-promoting cSCC milieu.

## INTRODUCTION

Epithelial tissues are composed of adherent sheets of polarized cells that serve important barrier functions. The epidermis is the body’s outermost and largest epithelial barrier, which is essential for both pathogen exclusion and fluid retention. The proper maintenance of epithelial integrity in this tissue relies on a resident stem cell population in the basal layer that ensures the rapid replacement of lost or damaged cells. These basal stem cells must balance between symmetric self-renewing divisions and production of non-proliferative suprabasal cells that are displaced upwards into overlying cell layers via asymmetric divisions^1^. These suprabasal layers will terminally differentiate to form the impenetrable skin barrier, and the outermost layers will die and be sloughed off. The skin undergoes constant turnover, where lost cells are replaced by additional basal stem cell asymmetric divisions that are fine-tuned to maintain proper tissue thickness.

Despite the importance of epithelial stem cells for skin tissue regeneration, their capacity to proliferate extensively as well as generate multiple cell types can also promote diverse disorders^2^. Notably, epithelial tissues are the source of 90% of human cancers, the most prevalent of which are cutaneous carcinomas, including squamous and basal cell carcinomas (cSCCs and BCCs, respectively), which primarily arise from epidermal basal stem cells^3^. Furthermore, aberrant stem cell behavior has been implicated in psoriasis, a chronic autoimmune skin condition where systemic inflammasome causes a buildup of scaly plaques^4^. Optimizing targeted treatments for skin disease requires a comprehensive understanding of the molecular players that promote stem cell dysregulation, which remains elusive. Our previous work revealed that genotoxic damage to homeostatic adult mouse skin results in activation of an innate immune signaling module called the inflammasome in dermal fibroblasts, which induces epithelial hyperplasia and stem cell fate misspecification via IL-1β signaling^5, 6^. Due to the potent impact of inflammasome activation on wildtype epidermal behavior, we hypothesized that inflammasome signaling may be involved in skin disease^7^. While the inflammasome pathway has been predominantly studied in the context of leukocyte-driven inflammatory processes, recent research has highlighted the potential role of this signaling to autoimmunity, metabolic and degenerative disease, and cancer^8–10^. Despite the conflicting correlative evidence that suggests a potential role for inflammasome signaling in skin cancer, mechanistic in vivo studies are lacking^11^. The inflammasome signaling pathway is canonically activated by damage- and pathogen-associated molecular patterns (DAMPs and PAMPs, respectively) that bind toll-like receptors (TLRs) on the cell surface, with the nucleotide-binding domain, leucine-rich–containing family, pyrin domain– containing-3 (NLRP3) inflammasome representing the predominant pathway known to be responsive to damage signals.

In this work, using both in vivo mouse models and human patient specimens, we discover that epidermal stem cell fate misspecification occurs across diverse pathophysiological settings, but not normal high proliferation contexts. We also reveal that inflammasome activation is not required for fate misspecification. Intriguingly, we determine that inflammasome activation is not generalizable to all epidermal disease but does occur during cSCC development. Based on our findings, we propose an cSCC tumor-promoting feed-forward signaling loop whereby kirsten rat sarcoma virus (KRAS)-induced inflammasome activation in epidermal epithelia promotes IL-1 signaling in dermal fibroblasts. This is a potentially potent cancer-driving mechanism that may be conserved across diverse epithelia.

## RESULTS

### Epidermal cell fate misspecification occurs in diverse pathophysiological contexts in mice

Our recent work revealed that epidermal stem cell misspecification occurs upon genotoxic damage^5^. We sought to determine whether this phenotype was a generalizable response to other skin aberrancies. To acquire more quantitative data than our prior work, we first recapitulated our previous findings by inducing epidermal hyperplasia in wildtype mouse backskin epidermis via topical application of the DNA crosslinking agent cisplatin and compared with vehicle-(PBS) treated or untouched skin as controls^5^. One week following treatment, we collected skin specimens and performed immunofluorescence staining and confocal imaging. We then applied a new quantitative approach to analyze stem cell fate misspecification by determining the percent thickness of epidermal layers co-expressing keratin 14 (K14), a basal stem cell marker, and K10, a suprabasal differentiation marker. Both untouched and vehicle-treated tissue exhibited a single K14-positive/K10-negative basal layer as well as K10-positive/K14-negative suprabasal layers, with very minimal overlapping K14/K10 double-positive expression evident in the bottommost suprabasal layer (Figure 1A). In cisplatin-treated tissue, however, we noted a significant increase in epidermal thickening concurrent with suprabasal cell fate misspecification, with the majority of suprabasal layers exhibiting double-positive K14/K10 expression, in contrast to the basal layer that maintained K14-positive/K10-negative expression (Figure 1A). These findings, which align with our previous work, suggest that cell fate misspecification occurs in epidermal suprabasal cells upon genotoxic insult, while also yielding a quantitative dataset with which we can compare to other aberrant skin states^5^.

**Figure 1.**
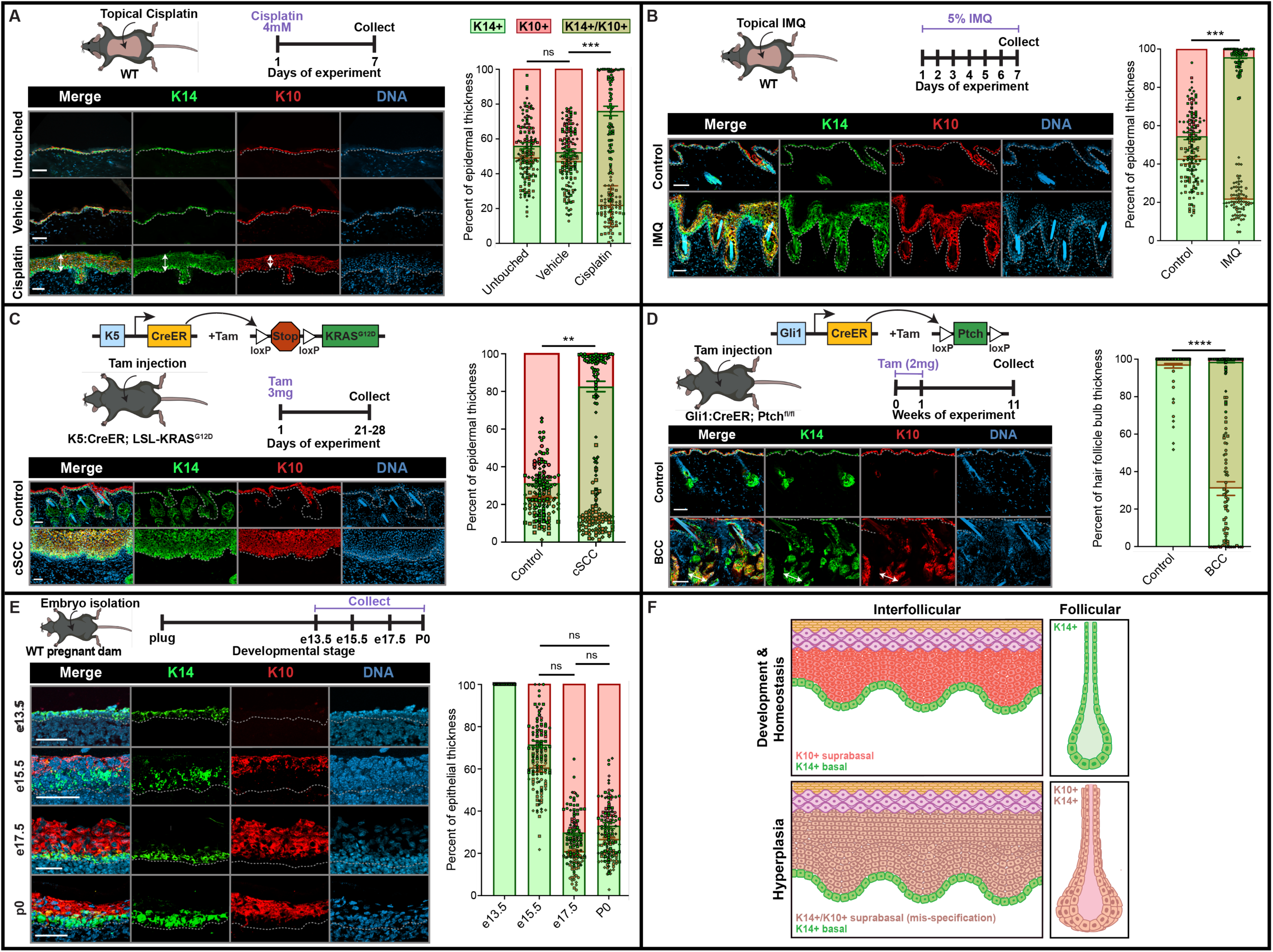
Epidermal cell fate misspecification occurs in diverse pathophysiological contexts in mice. **A-B)** Schematic of treatment and experimental timeline for inducing DNA damage (A) or psoriasis (B) in wildtype mouse skin, accompanied by corresponding immunofluorescence images of tissue 1-week post-treatment with topical 4mM cisplatin vs. PBS vehicle (A) or 5% imiquimod (IMQ) cream vs. shaved/untreated control (B). **C-D)** Schematic of transgenic construct, treatment, and experimental timeline for generating cutaneous squamous cell carcinoma (cSCC) (C) or basal cell carcinoma (BCC) (D) tumors in mice, accompanied by corresponding immunofluorescence images of tamoxifen-injected tumor and control backskin. **E)** Schematic and timeline of pup isolation at varying developmental stages from wildtype pregnant dams, accompanied by corresponding immunofluorescence images of embryonic (e) and postnatal day 0 (P0) skin. **A-E)** Each set of immunofluorescence images shows labeling of interfollicular epidermal stem cell marker keratin (K) 14 (green), differentiation marker K10 (red), and DNA (blue). Each dataset is accompanied on the right by bar graphs with scatter dot SuperPlots^57^ displaying the percent of K14-only (red), K10-only (green) and overlapping K14/K10 double-positive (olive) epithelial layers in interfollicular regions of A-C and E (see white arrows in A) or follicular bulb regions of D (see white arrows in D). Error bars represent standard error of the mean. N = 3 mice per bar. Individual values within each n are plotted in the same shape (circle, diamond, or square) and colored based on whether the measurement is K14 (green) or K10 (red). P-values, p > 0.05 (ns, not significant), p ≤ 0.01 (**), p ≤ 0.001 (***), p ≤ 0.0001 (****), were calculated using ordinary one-way analysis of variance (ANOVA) with Tukey’s multiple comparison tests (A and E) or unpaired student’s *t*-tests (B-D). Significance in A-C and E (where interfollicular epidermal regions were measured) was calculated based on K14 data, whereas D (where hair follicle bulb regions were measured) was determined based on K10 data. Dotted lines demarcate location of basement membrane. Scale bars = 50 µm. **F)** Schematic depicting normal stem cell specification during epidermal development and homeostasis, and aberrant stem cell specification (defined by overlapping suprabasal K14 and K10 expression) in hyperplastic skin contexts.

To determine whether a similar misspecification phenotype occurs in other epidermal hyperproliferative contexts or is a specific response to DNA crosslinking, we induced psoriasis, a chronic autoimmune disorder, in adult mouse backskin via topical application of the TLR7 agonist imiquimod (IMQ) using a well-validated protocol (5% IMQ daily for one week)^12^ (Figure 1B). Compared with controls, IMQ-treated skin showed hyper-thickening as well as a significant increase in overlapping K14/K10 expression in the suprabasal layers while the basal layer maintained K14-only expression, similar to Cisplatin-treated skin (Figure 1B).

We further extended these analyses to tumorigenic contexts in mouse skin by applying validated transgenic models to determine whether cutaneous cancers also result in epithelial fate misspecification. To generate cutaneous squamous cell carcinoma (cSCC), we induced constitutive expression of hyperactivated KRAS^G12D^ in the epidermis by injecting tamoxifen into K5: CreER; lox-STOP-lox (LSL)-KRAS^G12D^ mice^13^ (Figure 1C). Between three- and four-weeks post-induction, we isolated tumors from KRAS^G12D^-activated lip skin as well as normal skin from controls (tamoxifen-injected, KRAS^G12D^-negative). Notably, this mouse model was previously shown to favor skin tumor formation around oral regions over other anatomical sites^13^. Consistent with our observations in cisplatin- and IMQ-treated tissue, cell fate misspecification was present throughout the expanded suprabasal layers of cSCC tumors but not tamoxifen-injected controls (Figure 1C).

To assess whether this phenotype occurs across distinct cutaneous cancer contexts, we induced basal cell carcinoma (BCC) by hyperactivating the Sonic Hedgehog signaling pathway in mouse hair follicles via deletion of the tumor suppressor protein Patched (Ptch), the most mutated gene in human BCCs^14^. Importantly, hair follicles (which express Gli1) are a common BCC site of origin^15^. To drive BCC, we injected tamoxifen into Gli1: CreER; Ptch^fl/fl^ mice and collected backskin 11 weeks post-induction. Compared to control hair follicle bulbs, which are K14-positive and K10-negative, BCC hair follicles showed significant overlapping K14/K10 expression, revealing that misspecification also occurs during BCC tumorigenesis (Figure 1D)^15^. This phenotype is particularly striking, as K10 is an interfollicular differentiation marker that is not expressed in wildtype hair follicles. Notably, a similar hair follicle misspecification phenotype was observed when Cisplatin was intradermally injected into wildtype mouse skin^5^. These data demonstrate that fate misspecification under hyperplastic conditions is a generalizable phenomenon across distinct epidermal epithelial sub-compartments.

Taken together, these findings underscore that epithelial stem cell fate misspecification occurs in various hyperplastic and pathophysiological contexts. This suggests that overlapping K14/K10 expression could serve as an effective early biomarker of diverse skin pathologies should this misspecification phenotype be conserved in human disease contexts.

### Human cutaneous skin cancers exhibit epithelial stem cell fate misspecification

To establish whether stem cell misspecification also occurs in cancerous contexts in human skin, we procured fresh patient cSCC and BCC tumor specimens excised by a Mohs micrographic surgeon. Patient-matched non-cancerous tissue flanking each tumor, termed “cones”, were also obtained. As additional controls, skin specimens were collected from excess tissue isolated from patients who underwent abdominoplasty or mammoplasty procedures. Each specimen was cryoembedded, sectioned, and analyzed for K14 and K10 expression by immunofluorescence. In human control, cSCC cone, and BCC cone tissue, K14 expression was primarily restricted to epidermal basal cells with no significant expression observed in the K10-positive suprabasal layers (Figure S1A-B). In contrast, while the basal layer maintained K14-only expression in cSCC and BCC tumors, the suprabasal layers in these specimens showed double positive K14/K10 expression resembling our findings in mouse tumor tissue (Figure S1A-B). These data suggest that suprabasal cell fate misspecification in diseased skin is conserved between mice and humans.

### Proper epidermal fate specification is preserved in proliferative physiological contexts

We next sought to understand whether cell fate misspecification occurs under all high proliferation circumstances or is specific to aberrant skin. During embryogenesis, extensive epidermal cell proliferation facilitates epithelial surface area expansion of the elongating embryo as well as multilayering to establish an intact skin barrier prior to birth^16^. We collected embryos from wildtype pregnant dams at embryonic days 13.5, 15.5, and 17.5 as well as wildtype neonates at postnatal day 0 (p0) (Figure 1E). We then cryoembedded, sectioned, and immunostained the epidermis at each developmental stage for K14 and K10. We did not observe any K10 expression in e13.5 epidermis, which was consistent with the presence of only a single layer of K14-positive epithelial basal cells at this timepoint (Figure 1E). Nevertheless, in all other analyzed developmental stages, we noted a clear distinction between the K14-only basal layer and K10-only suprabasal layers with minimally overlapping K14/K10 expression at the basal/suprabasal interface (Figure 1E), comparable with our observations in untouched wildtype adult epidermis (Figure 1A). Taken together, these data suggest that stem cell fate misspecification is a characteristic of aberrant hyperplastic skin conditions, and that cell specification integrity is sustained during proliferative physiological contexts such as skin morphogenesis (Figure 1F).

### Inflammasome activation occurs in cSCC tumors

Our prior studies revealed that inflammasome activation drives hyperplasia and stem cell fate aberrancies in normal skin following genotoxic insult^5^. Inflammasomes are supramolecular organizing centers that catalyze IL-1β secretion in response to damage signals and have primarily been associated with innate immune responses (Figure 2A). Nevertheless, our work and others revealed that inflammasome activity serves important functions in non-immune cells, including epidermal epithelial cells and dermal fibroblasts^5, 17^. Considering our discovery that epidermal stem cell misspecification arises in diverse hyperplastic contexts, including cancer (Figures 1C-D, S1), we hypothesized that inflammasome activation may occur during skin tumorigenesis. Using the previously described KRAS^G12D^ hyperactivation mouse model, we generated cSCCs tumors in mice^13, 18–20^ and performed immunostaining for the core inflammasome component ASC (Figure 2A), which is known to aggregate into discrete “specks” upon inflammasome activation^21, 22^. We observed ASC specks in both the epithelial and dermal compartments of cSCC tumor tissue, while control tissue was devoid of specks (Figure 2B-C). Notably, we did not detect ASC specks in embryonic skin, psoriatic skin, or BCC tumors, suggesting that inflammasome activation is not generalizable to all aberrant or high proliferative skin contexts and that stem cell misspecification can occur independently of inflammasome activation (Figure S2).

**Figure 2.**
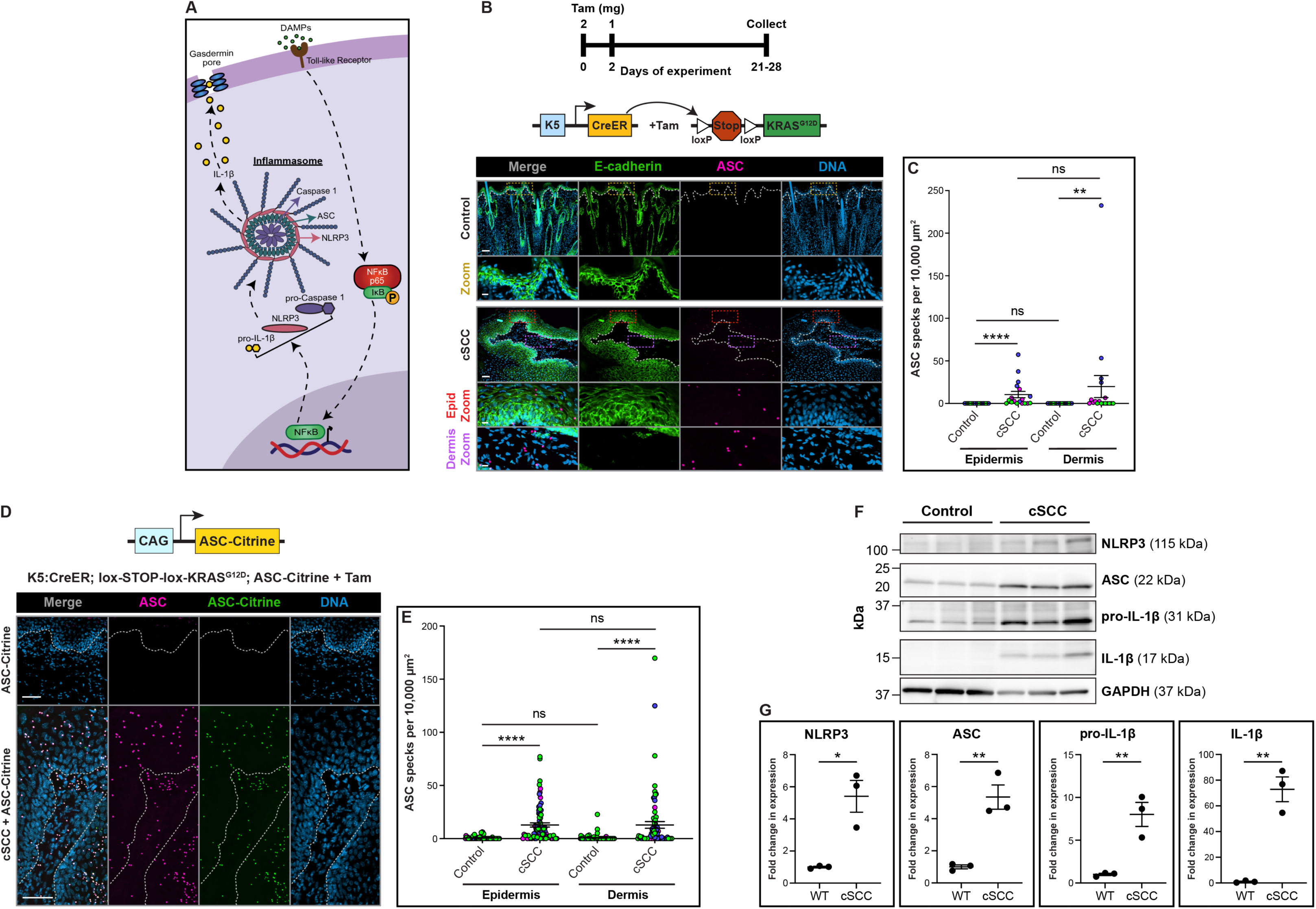
Inflammasome activation occurs in cutaneous squamous cell carcinoma (cSCC). **A)** Simplified schematic of NLR family pyrin domain containing 3 (NLRP3) inflammasome pathway. **B)** Experimental timeline for generating cSCC tumors in mice via tamoxifen (tam) induction accompanied by diagram of transgenic constructs. Representative immunofluorescence images show epithelial marker E-cadherin, inflammasome marker apoptosis-associated speck-like (ASC) protein (pink), which aggregates into discrete specks upon inflammasome activation, and DNA (blue). Dotted lines demarcate location of basement membrane. Colored dashed boxes correspond to epidermal (epid, red) and dermal (purple) zoom-in images. Scale bars = 50 µm. Epid, epidermis; Tam, tamoxifen; ER, estrogen receptor. **C)** Scatter dot SuperPlots displaying quantification of ASC specks in epidermal and dermal compartments of mouse cSCC tumors compared to control backskin. Error bars represent standard error of the mean. N = 3 mice per dot plot. Individual values within each n are plotted in the same color. P-values, p > 0.05 (ns, not significant), p ≤ 0.01 (**), p ≤ 0.0001 (****), were calculated using ordinary one-way ANOVA with Tukey’s multiple comparison tests. **D)** Diagram of ASC-citrine transgenic construct accompanied by representative immunofluorescence images of ASC immunostaining (pink) and citrine signal (green) in normal (ASC-citrine only + tamoxifen (tam)) or cSCC (K5: CreER; LSL-KRAS^G12D^; ASC-citrine + tam) backskin. Dotted lines demarcate location of basement membrane. Scale bars = 50 µm. **E)** Scatter dot SuperPlots displaying quantification of ASC specks (defined by specks positive for both citrine and ASC) in epidermal and dermal compartments of mouse cSCC tumors compared to control backskin. Error bars represent standard error of the mean. N = 3 mice per dot plot. Individual values within each n are plotted in the same color. P-values, p > 0.05 (ns, not significant), p ≤ 0.0001 (****), were calculated using ordinary one-way ANOVA with Tukey’s multiple comparison tests. **F)** Western blots showing inflammasome protein expression in mouse control and cSCC backskin. Each lane represents a different mouse (N = 3). Glyceraldehyde 3-phosphate dehydrogenase (GAPDH) is included as a loading control. kDa, kilodalton. **G)** Dot plots showing quantification of western blot in (F) with each protein signal normalized to its corresponding GAPDH signal. P-values, p < 0.05 (*), p ≤ 0.01 (**), were calculated using unpaired student’s *t*-tests.

To validate that the observed ASC puncta in cSCC tumor tissue are bona fide specks, we crossed an R26-CAG-LSL-ASC-citrine (“ASC-citrine”) line^23^ with K5:CreER; LSL-KRAS^G12D^ mice and induced tumorigenesis via tamoxifen injection. We observed a significant increase in ASC-citrine specks in cSCC tumors versus control (ASC-Citrine positive, KRAS^G12D^-negative) tissue, which was apparent in both the epidermal and dermal compartments (Figure 2D-E). Furthermore, we performed ASC immunostaining in K5: CreER; LSL-KRAS^G12D^; ASC-citrine tumors and confirmed endogenous ASC and ASC-citrine speck colocalization (Figure 2D)^23^. While a minimal number of ASC-citrine specks were present in control skin, this is consistent with previous reports that ASC overexpression, which was upregulated over 8-fold in the epidermis and over 11-fold in the dermis of ASC-citrine mice compared to wildtype skin (Figure S3A-B), induces spontaneous speck formation^24^. To further confirm inflammasome activation, we isolated control and cSCC tumor tissue from mouse skin and performed immunoblotting for the inflammasome components NLRP3, ASC, and IL-1β. Compared to control, the expression levels of all inflammasome proteins were significantly upregulated in cSCC tumors, including the processed form of IL-1β, which must undergo inflammasome-catalyzed cleavage prior to secretion (Figure 2F-G). We corroborated these data by determining that these same proteins are upregulated in human cSCC tumor tissue compared with control cone tissue (Figure S3C). Taken together, these data suggest that inflammasome activation occurs in both epithelial and dermal compartments of the skin during cSCC tumorigenesis.

### Inflammasome activation in KRAS-hyperactivated tissue precedes tumor formation and is epithelial autonomous

To gain deeper mechanistic insight into inflammasome activation within each cSCC skin compartment, we first tested signaling epistasis by exploring whether epithelial KRAS^G12D^ expression was sufficient to drive epithelial inflammasome signaling or whether dermal cues were required (Figure 3A). To do this, we isolated backskin from uninduced K5:CreER; LSL-KRAS^G12D^ mice, separated the epidermis from the dermis, plated the epithelium on laminin-coated culture dishes, and incubated the tissue in media supplemented with 4-hydroxytamoxifen (4-OHT) to induce CreER nuclear localization and subsequent KRAS^G12D^ expression (Figure 3B). Following ten days in culture, ASC specks were observed in KRAS^G12D^-expressing epithelium but not wildtype epithelium subjected to the same 4-OHT treatment (Figure 3C-D). These findings underscore that inflammasome activation in KRAS^G12D^-activated epithelium is epithelial autonomous. Importantly, the presence of ASC specks in cultured epithelium within days following KRAS ^G12D^ activation suggests that inflammasome activity precedes tumor formation.

**Figure 3.**
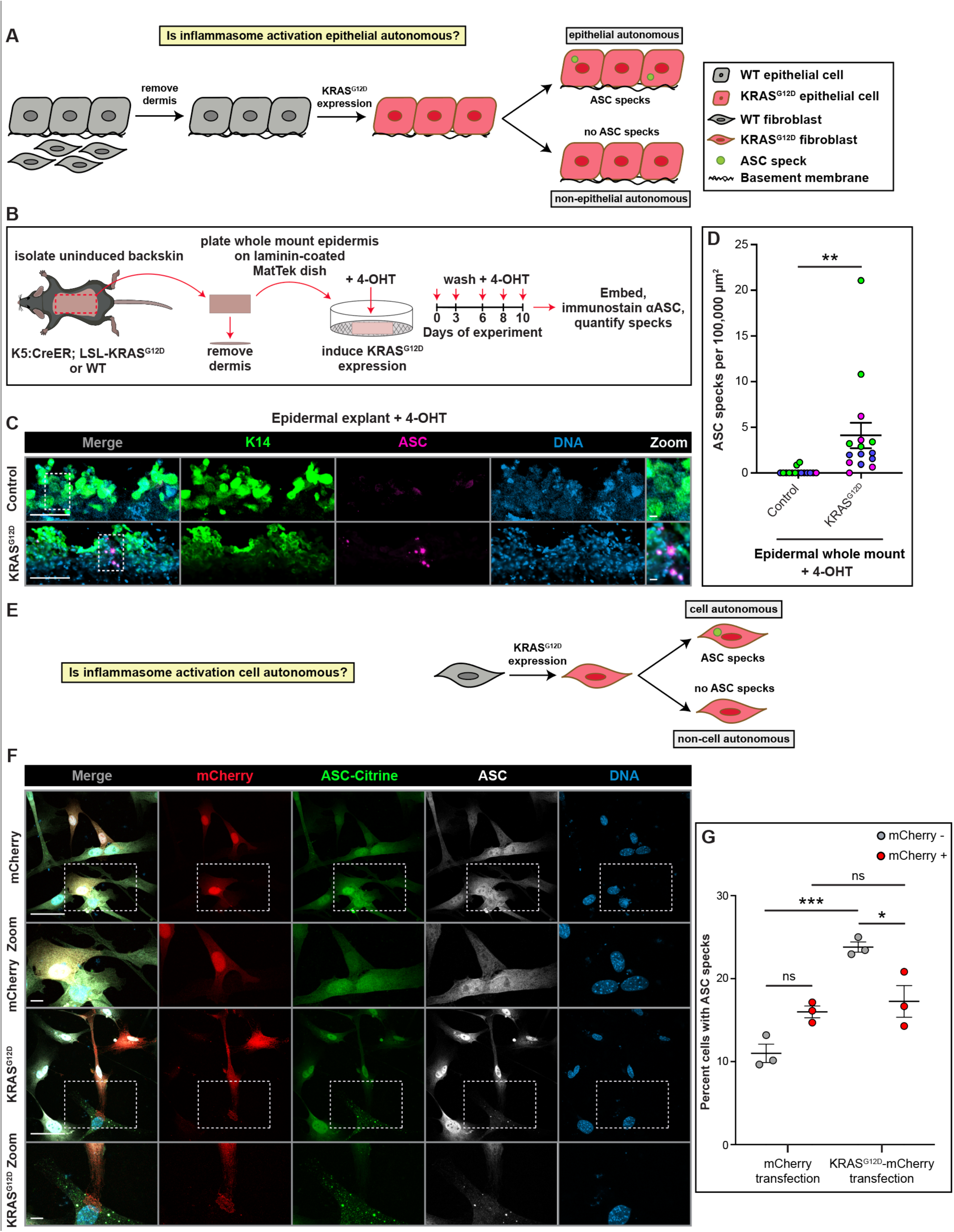
KRAS-driven inflammasome activation is epithelial autonomous but non-cell autonomous. **A)** Schematic of experimental results that would differentiate between epithelial autonomous versus non-autonomous inflammasome activation using an epidermal whole mount approach. **B)** Schematic of experimental approach to test whether KRAS-driven inflammasome activation is epithelial autonomous. **C)** Representative immunofluorescence images of ex vivo cultured KRAS^G12D^-expressing epidermal epithelium (isolated from K5: CreER; LSL-KRAS^G12D^ mice) containing ASC specks (pink) and control epithelium devoid of ASC specks (isolated from wild-type mice). Tissue was processed as outlined in (B). Scale bars = 50 µm (non-zoom) or 10 µm (zoom). **D)** Scatter dot SuperPlots displaying quantification of ASC specks in whole mount KRAS^G12D^-expressing and control epidermis. Error bars represent standard error of the mean. N = 3 mice per dot plot. Individual values within each n are plotted in the same color. P-value, p ≤ 0.01 (**), was calculated using unpaired student’s *t*-test. **E)** Schematic of experimental results that would differentiate between cell autonomous versus non-cell autonomous inflammasome activation using primary dermal fibroblasts. **F)** Representative immunofluorescence images of primary ASC-citrine fibroblasts transiently transfected with mCherry or KRAS^G12D^-mCherry plasmids. ASC immunostaining is shown in white. Scale bars = 50 µm (non-zoom) or 10 µm (zoom). **G)** Scatter dot plots displaying quantification of ASC specks (defined by specks positive for both citrine and ASC) in primary ASC-citrine dermal fibroblasts expressing control mCherry or KRAS^G12D^-mCherry compared to untransformed mCherry-negative fibroblasts present in the same transfection plates. Error bars represent standard error of the mean. N = 3 mice per dot plot (fibroblasts were freshly isolated from individual ASC-citrine mice). P-values, p > 0.05 (ns, not significant), p < 0.05 (*), p ≤ 0.001 (***), were calculated using ordinary one-way ANOVA with Tukey’s multiple comparison tests.

### Inflammasome activation in KRAS-hyperactivated tissue is non-cell autonomous

Considering KRAS is a component of the pro-proliferative mitogen-activated protein kinase (MAPK) signaling pathway, we sought to determine whether inflammasome activation occurs downstream of MAPK signaling in a cell autonomous manner following epithelial KRAS^G12D^ expression. Notably, published work has suggested a role for MAPK signaling in inflammasome priming and activation^25^. To rigorously test cell autonomous inflammasome activation, it was important to analyze a non-adhesive cell type to ensure our findings were interpretable and conclusive. We thus decided to freshly isolate primary dermal fibroblasts from wildtype ASC-citrine mice and transiently transfect these cells in culture with mCherry-KRAS^G12D^ (Figure 3E). As controls, we transfected ASC-citrine fibroblasts with mCherry alone. Notably, no significant differences in ASC speck numbers were detected when comparing 1) mCherry-positive versus negative cells in the mCherry-only transfection plates, or 2) mCherry-positive cells in the mCherry-only versus KRAS^G12D^-mCherry plates (Figure 3F-G). Strikingly, however, KRAS^G12D^-mCherry-positive cells had significantly fewer ASC specks than their neighboring untransfected cells (Figure 3F-G). Moreover, mCherry-negative cells in the KRAS^G12D^-mCherry transfection plates had significantly more ASC specks than mCherry-negative cells in the mCherry-only transfection plates (Figure 3F-G). These intriguing results suggest that KRAS^G12D^ hyperactivation in individual cells that lack intercellular connections is not sufficient to intrinsically activate the inflammasome but does induce sufficient DAMP production to activate inflammasomes in neighboring cells.

To confirm that primary fibroblasts can exhibit inflammasome activation in response to canonical signals, we treated freshly isolated ASC-citrine dermal fibroblasts with a combination of adenosine triphosphate (ATP), an established DAMP, and lipopolysaccharide (LPS), an established PAMP, which are known to activate inflammasomes in other cells types^26^. The combination ATP + LPS treatment, but not vehicle or LPS alone, was sufficient to induce ASC speck formation (Figure S3D). Our cumulative data suggest that KRAS^G12D^ hyperactivation in epithelia promotes non-cell autonomous inflammasome activation in nearby epithelial and dermal cells, likely via epithelial-derived DAMPs that bind transmembrane TLRs on these neighboring cell populations.

### Expansion, accumulation, and IL-1 signaling upregulation in cSCC fibroblasts

Considering our previously published findings of inflammasome-mediated fibroblast/epithelial crosstalk in hyperplastic skin^5^, we hypothesized that close fibroblast proximity to inflammasome-activated cSCC epithelial cells could facilitate epithelial-to-dermal IL-1β signal transduction. To visualize fibroblasts in normal and cSCC dermis, we immunostained mouse skin with two distinct fibroblast antibodies, Pdgfrα and vimentin, as well as the promiscuous leukocyte marker CD11b for comparison. Whereas immune cells remained scattered throughout the dermis, we noted significant fibroblast expansion and accumulation at the epithelial interface of cSCC tissue compared to control (Figure 4A).

**Figure 4.**
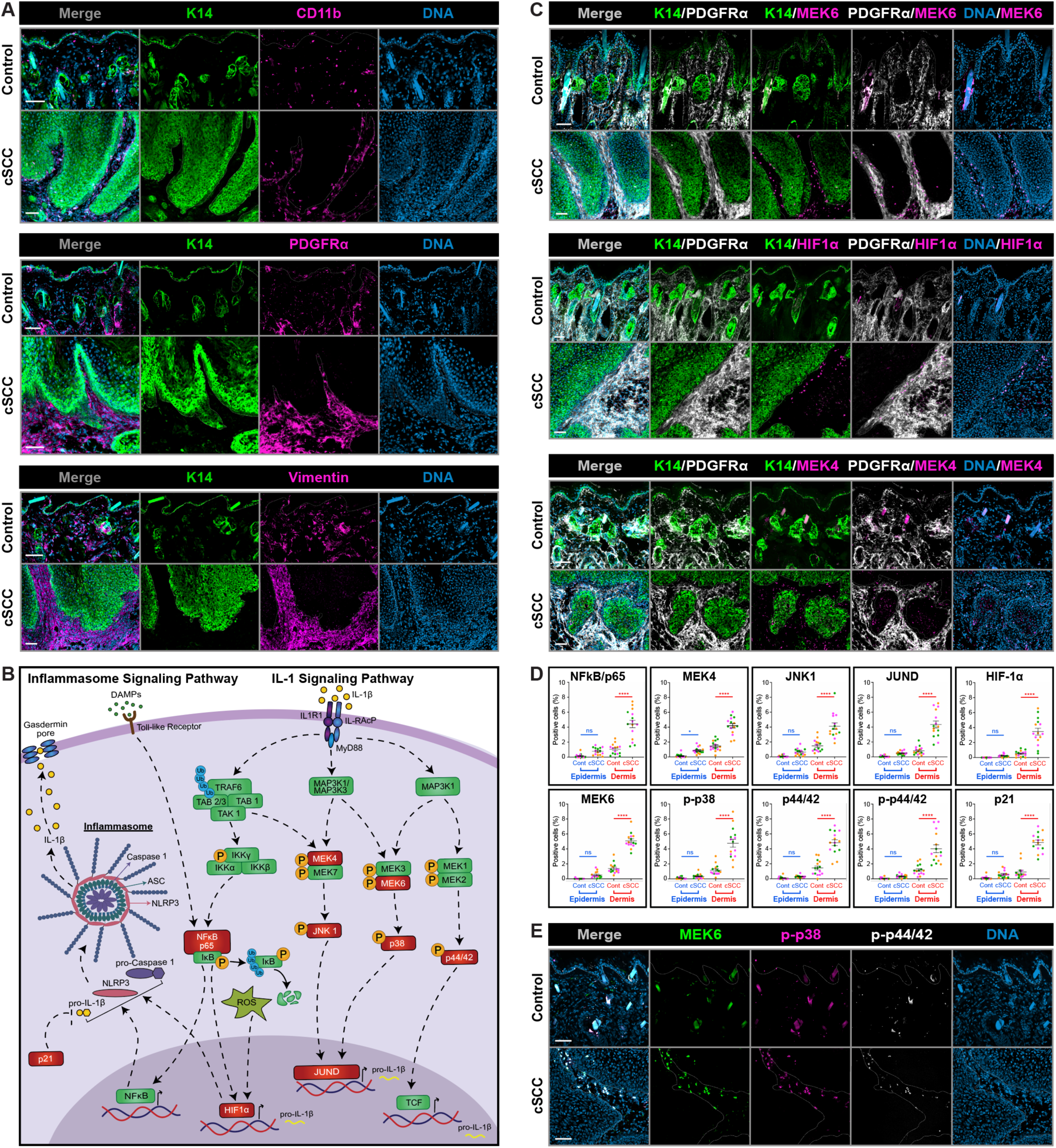
Expansion, accumulation, and IL-1 signaling upregulation in mouse cSCC fibroblasts. **A)** Representative immunofluorescence images of control and cSCC mouse tissue stained for epidermal stem cell marker K14 (green), leukocyte marker CD11b (pink), and fibroblasts markers Pdgfrα and vimentin (pink). Dotted lines demarcate location of basement membrane. Scale bars = 50 µm. **B)** Schematic of inflammasome and IL-1 signaling pathways. Proteins in red are analyzed in (C-E). **C)** Representative immunofluorescence images of control and cSCC mouse tissue stained for epidermal stem cell marker K14 (green), fibroblast marker Pdgfrα (white), as well as the IL-1 signaling proteins MEK6, Hif1α, and MEK4 (pink). Dotted lines demarcate location of basement membrane. Scale bars = 50 µm. **D)** Scatter dot SuperPlots displaying the percent of epithelial and dermal cells staining positive for IL-1 signaling proteins (labeled in red in (A)) in control and cSCC mouse tissue. Error bars represent standard error of the mean. N = 3 mice per dot plot. Individual values within each n are plotted in the same color. P-values, p > 0.05 (ns, not significant), p < 0.05 (*), p ≤ 0.0001 (****), were calculated using unpaired student’s *t*-tests. **E)** Control and cSCC mouse tissue co-stained for multiple IL-1 signaling components, confirming the same fibroblast subpopulations co-express all three targets. Dotted lines demarcate location of basement membrane. Scale bars = 50 µm.

To determine whether fibroblasts in cSCC skin likely respond to secreted IL-1β, we co-immunostained mouse control and cSCC tissue with Pdgfrα and the epithelial marker K14 along with a panel of antibodies targeting IL-1 signaling pathway proteins (see red colored genes in Figure 4B). We visualized a minimal population of IL-1 signaling-positive cells in the epidermal and dermal compartments of control skin, whereas cSCC tissue showed a significant increase in Pdgfrα-positive dermal but not epidermal labeling for all analyzed targets (Figure 4C-D). To confirm that these IL-1 signaling proteins were co-expressed in the same dermal fibroblast subpopulation, we performed co-immunostaining with antibodies against the IL-1 pathway proteins MEK6, phospho-p38MAPK, and phospho-p44/42 MAPK and noted triple-positive expression in the same cells (Figure 3E).

Integrating these data with our inflammasome activation findings (Figure 2), we propose a working model whereby KRAS hyperactivation induces epithelial DAMP release and binding to epithelial and dermal TLRs, which promotes non-cell autonomous inflammasome activation (Figure 5). Inflammasome signaling leads to IL-1β secretion from these cell populations, followed by preferential binding of IL-1β to IL-1 receptors on dermal fibroblasts (Figure 5). Notably, the finding that IL-1 signaling is absent from cSCC epithelia implies that this cell population may express insufficient levels of IL-1 membrane receptors. Importantly, the NFκB pathway is known to act downstream of both TLR and IL-1 receptor signaling, therein promoting an autocrine and/or juxtacrine IL-1β/IL-1 signaling feed-forward loop among neighboring fibroblasts to facilitate cSCC tumor growth (Figure 5)^27^.

**Figure 5.**
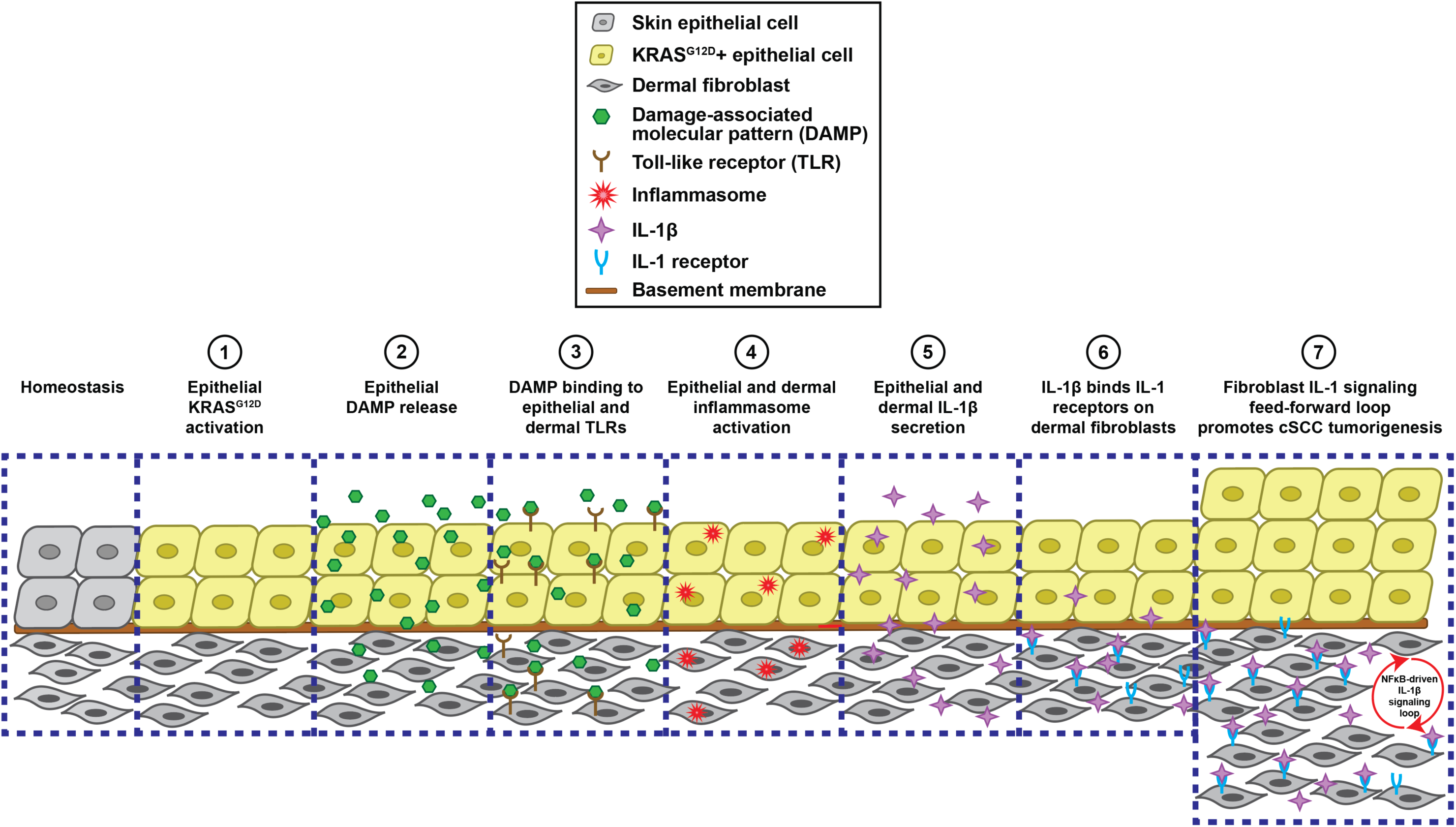
Working model of cSCC inflammasome activation and IL-1 feed-forward signaling. Cartoon depicting each step of the proposed mechanism suggested by our data, beginning with epidermal epithelial KRAS hyperactivation that ultimately leads to fibroblast IL-1 feed-forward signaling, which we suggest contributes to cSCC tumorigenesis.

## DISCUSSION

Taken together, our findings reveal that human and mouse skin stem cells exhibit defects in lineage specification in diverse disease contexts compared to normal skin, phenotypes that we previously observed in skin exposed to genotoxic agents^5, 7^. Specifically, suprabasal cells that are normally K10-positive and K14-negative instead co-express K10 and K14^28^. Importantly, K14 overexpression has been implicated in driving squamous cell carcinomas in diverse epithelia, including ovaries^29^, lung^30^, and skin^19^, and recent studies have identified important regulators of basal-to-suprabasal transitioning that prevent aberrant stem cell expansion^31^. Furthermore, while K10 expression has been shown to inhibit skin proliferation and tumorigenesis^32^, K10-positive suprabasal cells can serve as cells of origin for cutaneous squamous cell carcinomas^19, 33^. These discoveries suggest that epidermal stem cell fate misspecification may serve as a reliable biomarker of skin disease. Notably, previous work proposes causal links between aberrant stem cell lineage specification and skin disorders, including cutaneous skin cancers^34^ and psoriasis^4^. Furthermore, lineage misspecification can promote intertumoral heterogeneity, causing cancers to become refractory to chemotherapy^35, 36^.

Our studies further demonstrate that inflammasome activation occurs in response to epithelial KRAS hyperactivation and that this likely initiates feed-forward IL-1 signaling in cSCC fibroblasts. Notably, IL-1 signaling feedback is known to drive cancer, degenerative disease, and autoinflammatory disorders in diverse tissues^37, 38^. The observed expansion, inflammasome activation, and IL-1 signaling in cSCC fibroblasts may suggest that these cells have transitioned into cancer-associated fibroblasts (CAFs), a classification linked to the progression of cutaneous carcinomas as well as many other diverse epithelial cancers^39–42^. Notably, pancreatic CAFs were recently reported to assume an inflammatory nature (“iCAFs”) in response to IL-1 signaling^43, 44^, and breast cancer CAFs can exhibit inflammasome activation^45^. Nevertheless, CAF-specific markers have not been clearly defined due to significant expression heterogeneity, as well as the considerable overlap among CAF, normal fibroblast, and immune cell protein signatures^46^. Additional studies are required to delineate the role of inflammasome and IL-1 signaling in the establishment of CAF identity and tumorigenicity.

The finding that inflammasome activation precedes tumor formation in KRAS^G12D^ epithelium suggests that inflammasome-driven IL-1 signaling is a cause and not an effect of skin tumorigenesis. Nevertheless, further research is needed to determine whether fibroblast feed-forward IL-1 signaling is necessary and/or sufficient for cSCC development, maintenance, and progression. It will also be important to determine the mechanism that underlies the observed heterogeneity in epidermal and dermal inflammasome activation as well as fibroblast IL-1 signaling during cSCC tumorigenesis. Whether these signaling subpopulations have greater tumorgenic, metastatic, and/or recurrent potential are interesting questions for future investigation.

## MATERIALS AND METHODS

### Lead Contact

Further information and requests for resources and reagents should be directed to and will be fulfilled by the lead contact, Lindsey Seldin.

### Materials Availability

Authors will share the mCherry-KRAS^G12D^ construct upon request.

### Animals

The mouse lines acquired from Jackson Laboratory (Bar Harbor, ME) for this study include: LSL-KRAS^G12D^ (JAX stock # 008180^47^), R26-CAG-LSL-ASC-citrine (Jax stock # 030743^23^) (referred to throughout text as “ASC-Citrine”), C57BL/6J (Jax stock # 000664^48^) and C3H (Jax stock # 000659^49^). K5:CreER mice (Jax stock # 029155^50^) were provided by Dr. Brigid Hogan from Duke University and Gli1:CreER^T2^; Ptch floxed mice^51^ (Jax stock # 007913^52^ and # 012457^53^, respectively) were provided by Dr. Sunny Wong from University of Michigan. Skin experiments were performed on 3-12 wk old adult mice, both males and females. Littermates (when possible) or age-matched animals were used for experimental and control treatments. Mice were allocated to experimental groups based on genotype (when applicable) or randomly. Mice were housed with a standard 12 h light/12 h dark cycle. Mice were provided normal laboratory chow and water. All mouse experiments were performed with approval from the Institutional Animal Care and Use Committees at both Emory University and the Atlanta VA Medical Center. Numbers of mice used for each experiment can be found in the corresponding figure legends.

### Disease Induction in Mice

*<u>DNA damage:</u>* Following isoflurane anesthesia administration via nose cone using a low-flow electronic vaporizer^54^, adult wildtype C3H mouse backskin was shaven and superficial abrasions were made using a Wahl razor (Wahl Clipper Corporation, Sterling, IL) prior to topical application of 4mM Cisplatin (BioVision, Inc., Milpitas, CA) or PBS (“vehicle”) or nothing (“untouched”)^5^. Mice were sacrificed and backskin samples were cryoembedded on day 7 post-treatment. *<u>Psoriasis:</u>* Adult wildtype C57BL/6J mice were anesthetized as previously described and 15 mg cream containing 5% IMQ (Taro Pharmaceutical Industries, Haifa, Israel) was topically applied to shaven backskin daily for 7 days^12^. Shaven, untreated mice were used as controls. Mice were sacrificed and backskin samples were cryoembedded on day 7 post-treatment. *<u>Tumorigenesis:</u>*To induce cSCC tumors, K5: CreER; LSL-KRAS^G12D^ ^fl/wt^ (+/- ASC-citrine) were intraperitoneally injected once with 3 mg of tamoxifen dissolved in corn oil (Aqua Solutions, Inc., Deer Park, TX). Mice were sacrificed and tumor-containing skin (consistently concentrated around the mouth) was collected and cryoembedded 4 wks post-injection. Skin specimens from control mice (tamoxifen-injected animals with no KRAS induction) were collected from the same anatomical region. Gli1: CreER; Ptch^fl/fl^ mice were intraperitoneally injected twice (on experimental days 1 and 7) with 2 mg of tamoxifen. Mice were sacrificed and backskin samples were cryoembedded 11 wks post-treatment. Control backskin was collected from tamoxifen-injected Gli1: CreER; Ptch^fl/wt^ mice, as these animals did not show signs of tumor growth.

### Mouse Embryo Collection

Wildtype C57BL/6J males and females were mated and females were checked daily for plugs. Upon plug detection, developmental staging was tracked, and pregnant dams were sacrificed for embryo collection at embryonic (e) days 13, 15 and 17. Surgical scissors were used to make an abdominal incision, and embryos were pulled from abdominal cavity, dislodged from their individual embryonic sacs using scissors, washed with PBS, and cryoembedded.

### Human Tissue Procurement

Patient skin tissue, including squamous and basal tumors as well as juxtaposed cone tissue, was isolated during routine Mohs surgery at the Emory Clinic. Excess tissue not required for pathology analyses was deidentified and provided to our lab the day of surgery suspended in Gibco RPMI 1640 Medium without Glutamine (ThermoFisher Scientific, Waltham, MA). Upon receipt, adipose tissue was removed using sterile surgical scissors, and the remaining tissue was washed in PBS and either cryoembedded for cryosectioning or added to lysis buffer for western blotting. This tissue acquisition was approved by the Emory and VA Institutional Review Boards (IRB) under ID # STUDY00004423 (non-human subjects). Excess control skin specimens isolated during routine abdominoplasty and mammoplasty procedures and deidentified were provided by the NCI-funded Cooperative Human Tissue Network (CHTN) Western Division located at Vanderbilt University Medical Center.

### Immunostaining

Primary antibodies used in this study included: chicken α-cytokeratin 14, rabbit α-cytokeratin 10, and rat α-E-cadherin (BioLegend, San Diego, CA), rabbit α-MEK-4, rabbit α-MEK-6, and rabbit α-JUND (Abcam, Cambridge, UK), rabbit α-IL-1β (ABclonal Technology, Woburn, MA), rat α-Pdgfrα (ThermoFisher Scientific, Waltham, MA), rat α-CD11b (BD Biosciences, San Jose, CA), rabbit α-NLRP3, rat α-vimentin, rabbit α-JNK1, and rabbit α-NFκB p65 (BioTechne, Minneapolis, MN), rabbit α-ASC (AdipoGen, San Diego, CA), rabbit α-HIF-1α, rabbit α-p44/42 MAPK, rabbit α-phospho-p44/42 MAPK, rabbit α-p21, rabbit α-phospho-p38 MAPK, and rabbit α-GAPDH (Cell Signaling Technology, Danvers, MA). Secondary antibodies included: Alexa Fluor 488, 594 and 647 conjugates of anti-chicken, -rat, and -rabbit (ThermoFisher Scientific, Waltham, MA). DNA was stained using Hoechst 33342 (ThermoFisher Scientific, Waltham, MA). The Zenon Tricolor Rabbit IgG Labeling Kit (ThermoFisher Scientific, Waltham, MA) was used to co-stain skin tissue with rabbit α-MEK6, rabbit α-phospho-p38, and rabbit α-phospho-p44/42 antibodies (Figure 4E).

For immunohistochemical analyses, mouse backskin specimens and epidermal explants were cryo-embedded in O.C.T. (Fisher Scientific, Hampton, NH). A Leica CM1850 cryostat was used to generate 12 µm tissue sections and mounted on ColorFrost Plus microscope slides (Fisher Scientific, Hampton, NH). Tissue and cell culture coverslips were fixed and permeabilized for 8-10 min in 4% paraformaldehyde (PFA) dissolved in phosphate buffered saline containing 0.2% Triton X-100 (PBST) and blocked for 30 min in a blocking buffer solution containing 0.2% PBST and 5% goat/donkey serum. Primary antibodies were diluted to their optimal concentration in blocking buffer solution and incubated on tissue for 15 minutes. Tissue or cells were then washed three times for 5 min each with 0.2% PBST and secondary antibodies (including Hoeschst DNA stain) were added for 10 min prior to an additional three rounds of 5 min washes with 0.2% PBST. Fluoromount G followed by #1.5 thickness cover glass (Electron Microscopy Sciences, Hatfield, PA) was added atop tissue and then sealed with clear nail polish. For cultured cells, a dollop of Fluoromount G was added to each stained coverslip, which was then laid cells-side-down onto glass microscope slides and sealed with clear nail polish.

### Confocal Microscopy

Immunofluorescence images were acquired using a Nikon AxR laser scanning confocal microscope, 60x Oil 1.42 numerical aperture (NA), 40xC Silicone 1.25 NA, and 20X Air 0.75 NA Plan Apochromat objectives, Type B Immersion oil and Nikon Silicone Immersion oil (Cargille Laboratories, Cedar Grove, NJ), and Nikon NIS-Elements imaging software. Fiji (ImageJ) software was utilized for post-acquisition processing. Images throughout manuscript are maximum intensity projections of z-stacks, and merged images are composites of individual color channels. Where noted in figure legends, images were taken using the same exposure times for each color channel.

### Mouse Dermal Fibroblast Isolation and Culture

Mouse neonatal pups were sacrificed by decapitation and the backskin was isolated using surgical scissors. Tissue was rinsed with sterile 70% ethanol and PBS and then incubated dermis side down overnight at 4°C in a 1:1 mixture of PBS and Dispase-II (Sigma-Aldrich, St. Louis, MO). The next day, the epidermis was pulled off the dermis using sterilized forceps and the dermis was minced with sterilized surgical scissors. Minced dermis was incubated in collagenase media (DMEM (Sigma Aldrich, St. Louis, MO) with high glucose supplemented with 1.25 mg/mL collagenase type I, 0.5 mg/mL collagenase type II, 0.5 mg/mL collagenase type IV, 50 U/mL DNase I (STEMCELL Technologies, Vancouver, Canada), and 0.1 mg/mL hyaluronidase IVS (Sigma Aldrich, St. Louis, MO)), for 20 mins at 37°C. Then, growth media (DMEM with high glucose supplemented with 10% FBS (GeminiBio, West Sacramento, CA), 100 U/mL penicillin, and 100 µg/ml streptomycin (Sigma Aldrich, St. Louis, MO)) was added to neutralize the enzymes. The mixture was poured through a 70 µm Falcon cell strainer followed by a 40 µm Falcon cell strainer (BD Biosciences, San Jose, CA) to collect fibroblasts. Cells were spun down at 1500 xg for 5 min, washed twice with growth media, and cultured in growth media at 37°C and 5% CO_2_.

### Mouse Dermal Fibroblast Transfection

Primary fibroblasts were isolated as described above from the backskin dermis of ASC-citrine neonatal mice and plated on 1.5 thickness cover glass (Electron Microscopy Sciences, Hatfield, PA) in 35 mm dishes in DMEM growth media (ingredients described in previous section). Control mCherry-only or experimental mCherry-KRAS^G12D^ plasmids were transiently transfected into these cells using Lipofectamine 3000 (ThermoFisher Scientific, Waltham, MA) according to the manufacturer’s instructions. In short, DMEM was replaced with Opti-Minimal Essential Medium (MEM) (ThermoFisher Scientific, Waltham, MA) immediately prior to transfection. A transfection mixture containing 2.5 µg of plasmid DNA was incubated with cells for 6 hours prior to removal and replacement with DMEM growth media. Cells were then cultured overnight at 37°C and 5% CO_2_ prior to fixation and immunostaining.

### Inflammasome Activation in Mouse Dermal Fibroblasts

Primary fibroblasts were isolated as described above from the backskin dermis of ASC-citrine neonatal mice. Freshly isolated fibroblasts were cultured for 24 hr and then treated with 1 µg/mL lipopolysaccharide (LPS) (Sigma Aldrich, St. Louis, MO) dissolved in PBS or PBS only (vehicle) for 24 hr. Then, 1-, 3-, or 5-mM ATP (Sigma Aldrich, St. Louis, MO) dissolved in water or water only (for vehicle and LPS only conditions) were added to the cells for 1 hr. Cells were then immunostained or collected to make protein lysates for western blotting.

### Whole Mount Epidermal Culture

Mouse neonatal backskin was isolated as described in the previous section. Following overnight incubation in 1:1 PBS and Dispase-II, the epidermis was pulled off the dermis using sterilized forceps, plated on a 35 mm glass-bottom dish (MatTek Corporation, Ashland, MA) coated with 100 µM Laminin (Sigma Aldrich, St. Louis, MO) and incubated for 3 hr at 37°C and 5% CO_2_ prior to addition of DMEM. Tissue was cultured in vehicle (100% EtOH) or 0.02 mg/mL 4-OH tamoxifen for 10 days prior to cryoembedding and immunostaining in late afternoon of the 10^th^ day. Fresh 4-OHT-supplemented media was added to tissue in the mornings of days 3, 6, 8, and 10 post-plating.

### Western Blotting

Cultured cell lysates were prepared using Triton X-100 lysis buffer (50 mM Tris-HCL, pH 7.4, 150 mM NaCl, 5 mM EDTA, and 1% Triton X-100) including M221 protease inhibitor (VWR, Radnor, PA). Tissue lysates including mouse epidermis and dermis as well as human skin specimens were homogenized by mincing and sonicating with ice-cold RIPA lysis buffer (50 mM Tris-HCL, pH 7.4, 150 mM NaCl, 5 mM EDTA, 0.1% SDS, 0.1% sodium deoxycholate, and 1% Triton X-100) with M221 protease inhibitor. Homogenates were centrifuged and supernatants were collected as the lysates. Protein concentrations for each sample were determined using the Pierce BCA Protein Assay Kit (ThermoFisher Scientific, Waltham, MA). Following separation with SDS-PAGE, proteins were transferred to a PVDF membrane using a wet-transfer system (Bio-Rad, Hercules, CA). Chemiluminescence signals were detected using the Radiance Plus reagent (Azure Biosystems, Dublin CA) and visualized using a ChemiDoc MP Imaging System (Bio-Rad, Hercules, CA).

### Cloning of mCherry-KRAS^G12D^ Construct

The pBabe-KrasG12D, pQCXIP-mCherry-Halo-YAP1, and mCherry2-C1 plasmids were acquired from Addgene (Watertown, MA). pBabe-Kras^G12D^ was a gift from Channing Der (Addgene plasmid # 58902; http://n2t.net/addgene:58902; RRID:Addgene_58902). The pQCXIP-mCherry-Halo-YAP1 plasmid was a gift from Yutaka Hata (Addgene plasmid # 128336; http://n2t.net/addgene:128336; RRID:Addgene_128336)^55^. mCherry2-C1 was a gift from Michael Davidson (Addgene plasmid # 54563; http://n2t.net/addgene:54563; RRID:Addgene_54563). The *Kras^G12D^* cDNA insert was PCR amplified from pBabe-Kras^G12D^ using the following primers: 5’ AAGCCCACGCGTACTGAATATAAACTTG and 3’ AATAAAGCGGCCGCTTACATAATTACACAC. The amplified product was digested with MluI-HF and NotI restriction enzymes (New England Biolabs, Ipswich, MA) and ligated into digested pQCXIP-mCherry-Halo-YAP1 plasmid, replacing the Yap1 sequence. The mCherry2-C1 plasmid was used as a control. All plasmids were sequence confirmed.

### Quantification and Statistical Analyses

*<u>Misspecification:</u>* Total epidermal thickness was determined using Fiji software by measuring from the bottommost K14 signal to the topmost K10 signal (see double-sided white arrows in Figure 1A). The thickness of the K14 layers and K10 layers were then measured separately in each respective channel, and their percentages were calculated based on total epidermal thickness. To determine the percent overlap, the K14 and K10 percentages were added together and subtracted from 100. Regions in or juxtaposed to hair follicles were excluded from these measurements. *<u>Western blotting</u>:* Protein bands in western blots were quantified using the densitometry feature in Fiji and normalized to the GAPDH loading control^56^. For ASC-Citrine western blots, the average of the lower value group in each graph was set at 1 and the larger values were normalized to 1 to determine fold-change. *<u>Graphs</u>:* Plots were generated using GraphPad Prism. Thick middle line in each plot represents standard error of the mean (SEM). Statistical analyses were performed using mixed model one-sided ANOVAs for multiple comparisons, and unpaired one-sided Student’s *t*-tests for single comparisons using GraphPad Prism software. Statistical significance was assigned to p-values ≤ 0.05 (p < 0.05 (*), p ≤ 0.01 (**), p ≤ 0.001 (***), p ≤ 0.0001 (****)). Statistical details of experiments can also be found in the figure legends (n refers to number of mice for in vivo experiments, and number of experimental replicates for ex vivo experiments). SuperPlots were generated using GraphPad Prism^57^.

## Acronyms

ASC: apoptosis-associated speck-like protein containing a CARD
ATP: adenosine triphosphate
BCA: Bicinchoninic acid
BCC: basal cell carcinoma
cSCC: cutaneous squamous cell carcinoma
CHTN: cooperative human tissue network
DAMP: damage-associated molecular pattern
DMEM: Dulbecco’s Modified Eagle Medium
EDTA: ethylenediaminetetraacetic acid
ER: estrogen receptor
GAPDH: glyceraldehyde 3 phosphate dehydrogenase
HCl: hydrochloric acid
IL-1β: interleukin 1 beta
IMQ: imiquimod
IRB: institutional review board
K10: keratin 10
K14: keratin 14
KRAS: kirsten rat sarcoma virus
LPS: lipopolysaccharide
MAPK: mitogen-activated protein kinase
MEK: mitogen-activated extracellular signal-regulated kinase
MEM: Minimal Essential Medium
NaCl: sodium chloride
NLRP3: nucleotide-binding domain, leucine-rich–containing family, pyrin domain–containing-3
NS: not significant
OCT: optimal cutting temperature
OHT: hydroxytamoxifen
PAMP: pathogen-associated molecular pattern
PBS: phosphate-buffered saline
Pdgfrα: platelet-derived growth factor receptor alpha Ptch, patched
PVDF: Polyvinylidene fluoride
RIPA: Radio-Immunoprecipitation Assay
RPMI: Roswell Park Memorial Institute
SDS-PAGE: sodium dodecyl-sulfate polyacrylamide gel electrophoresis
SEM: standard error of the mean
TLR: toll-like receptor

## ACKNOWLEDGEMENTS

We thank Ian Macara for reagents, Sunny Wong for the Gli1:CreER^T2^; Ptch floxed mice, and Brigid Hogan for the K5: CreER mice. This work was supported by Career Development Award IK2 BX005370 from the U.S. Department of Veterans Affairs Biomedical Laboratory R&D Service to LS. This manuscript is dedicated to the brilliant scientist Dr. Sriram Srikant, who recently lost his battle with cancer but whose legacy will be a continual source of inspiration.

## AUTHOR CONTRIBUTIONS

Conceptualization, LS; Methodology, LS, CL; Validation, LS; Formal Analysis, KRP, RS, KT, RW, MH, ABP, DS, LS; Investigation, KRP, RS, KT, RW, MH, NS, ABP, DS, JL, CES, LS; Data Curation, KRP, RS, KT, RW, MH, LS; Resources, AH, TB, CL; Writing - Original Draft, LS; Writing - Review and Editing, KRP, RS, RW, MH, LS; Visualization, KRP, RS, KT, RW, MH, NS, ABP, DS, JL, CES, LS; Supervision, LS; Funding Acquisition, LS.

## SUPPLEMENTARY FIGURE LEGENDS

**Figure S1.**
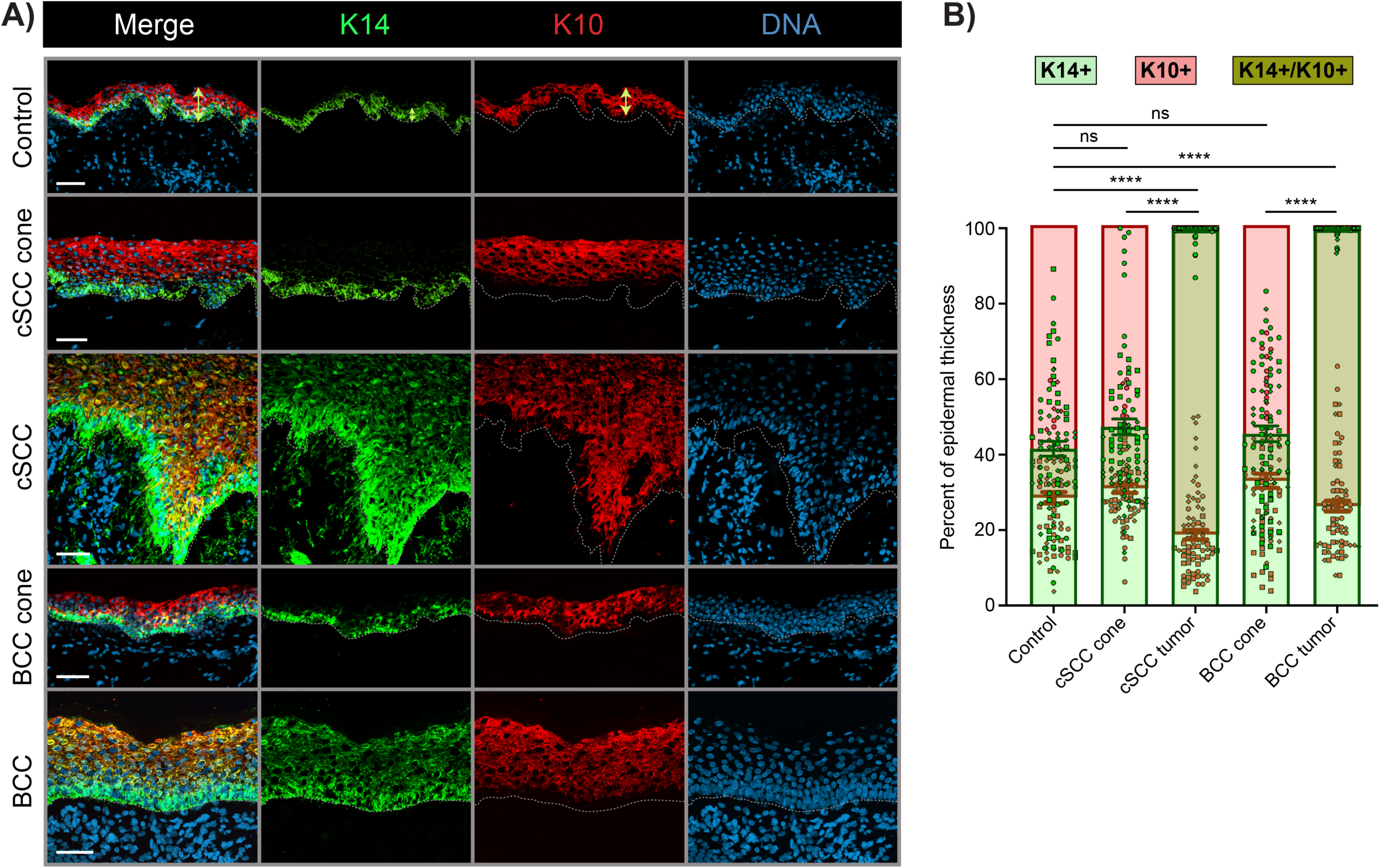
Human cutaneous skin cancers exhibit epithelial stem cell fate misspecification. **A)** Representative immunofluorescence images of human control, cone, and tumor epidermal tissue stained for epidermal basal stem cell marker K14 (green) and differentiation marker K10 (red) and DNA (blue). cSCC, cutaneous squamous cell carcinoma; BCC, basal cell carcinoma. “Control” refers to excess skin tissue acquired during abdominoplasties or reduction mammoplasties in non-cancerous patients; “Cone” refers to patient-matched skin juxtaposed to tumor isolated during Mohs surgery. Dotted lines mark location of basement membrane. Scale bars, 50 µm. **B)** Bar graphs with scatter dot SuperPlots displaying the percent of K14-only (red), K10-only (green) and overlapping K14/K10 double-positive (olive) epidermal epithelial layers with respect to total tissue thickness (see yellow arrows in Control panels of A). Error bars represent standard error of the mean. N = 3 human specimens per bar. Individual values within each n are plotted in the same shape (circle, diamond, or square). P-values, p > 0.05 (ns, not significant), p ≤ 0.0001 (****), were calculated using ordinary one-way ANOVA (based on K14/K10 overlap data) with Tukey’s multiple comparison test.

**Figure S2.**
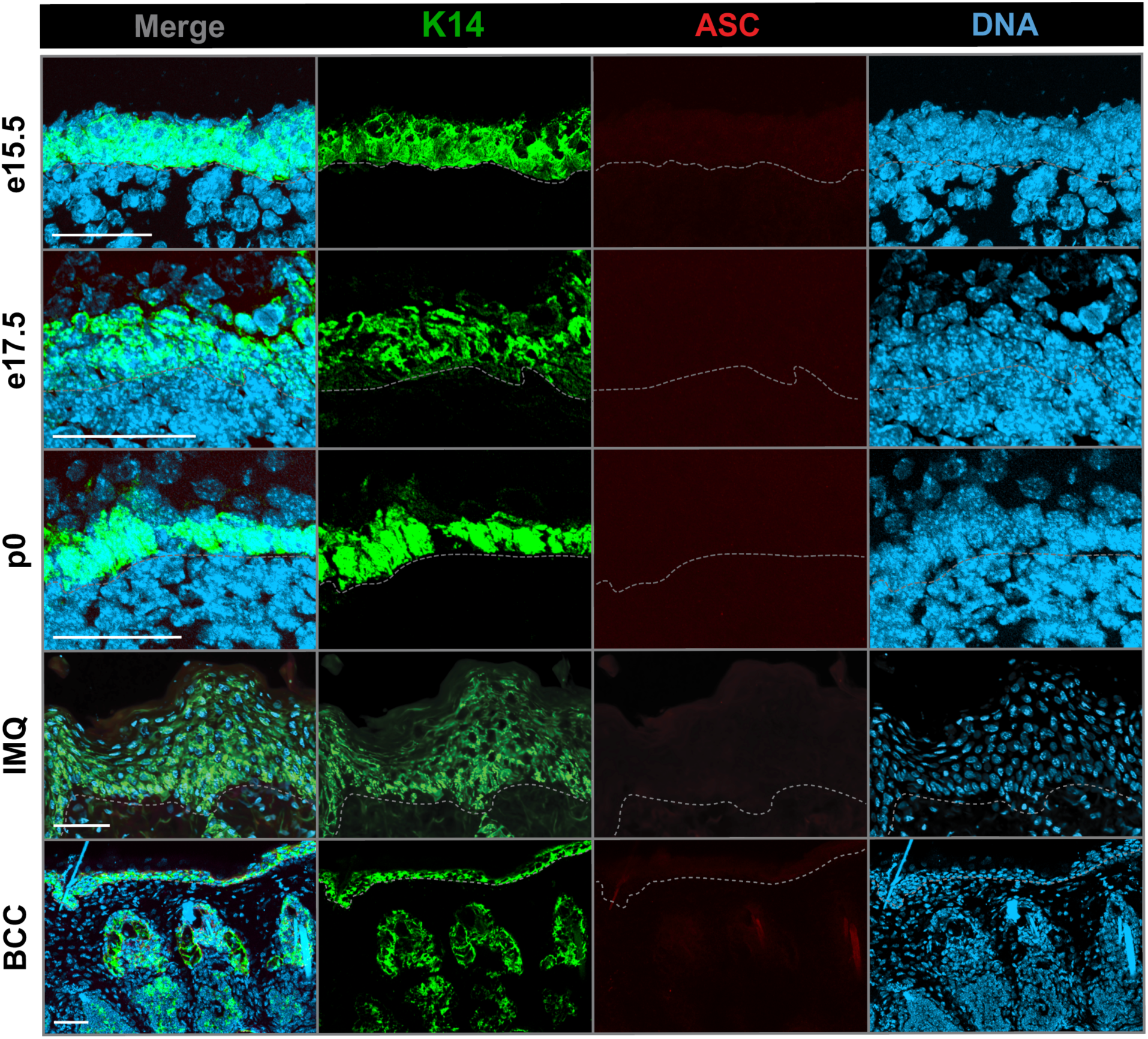
Inflammasome activation does not occur in mouse embryogenesis, psoriasis, or basal cell carcinoma. Representative immunofluorescence images of the epithelial stem cell marker K14 (green), the inflammasome marker ASC (pink) and DNA (blue) in mouse embryonic (e) and postnatal day 0 (P0) skin as well as imiquimod-treated backskin and basal cell carcinoma (BCC) tumors. Inflammasome activation was not detected in any of these contexts, evidenced by the absence of ASC specks. N = 3 mice analyzed per category. Quantification graphs are omitted due to zero ASC specks counted for each tissue sample.

**Figure S3.**
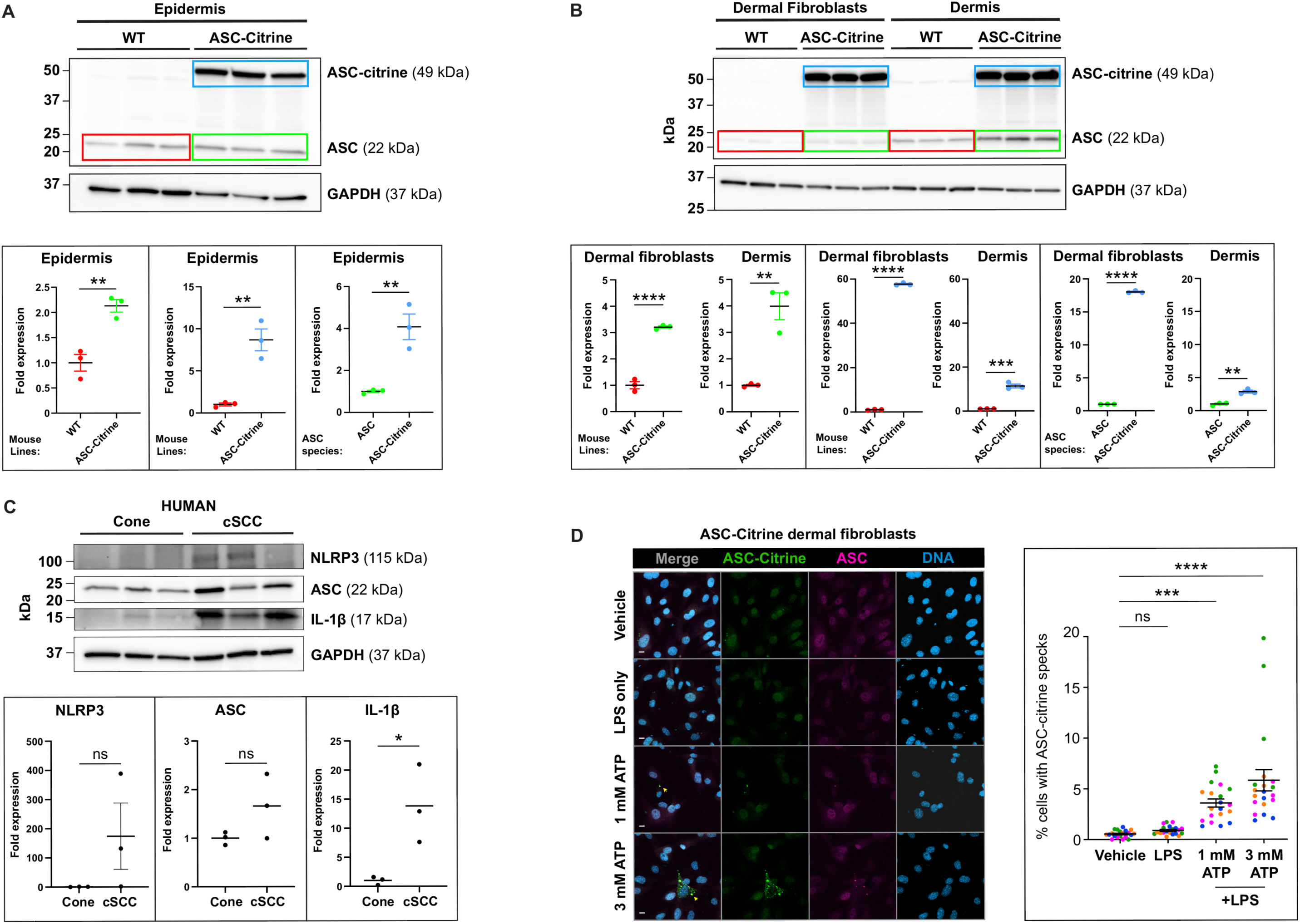
Inflammasome activation in mouse and human skin cells. **A-B)** Western blot comparing ASC protein levels in wild-type (WT) versus ASC-Citrine mouse epidermis (A) as well as dermis and dermal fibroblasts (B). Dot plots below each respective western compares the protein levels of endogenous ASC in wild-type tissue (red data and box), endogenous ASC in ASC-citrine tissue (green data and box), and ASC-citrine in ASC-citrine tissue (blue data and box). Each lane represents a different mouse (N = 3). kDa, kilodalton. **C)** Western blot comparing inflammasome protein levels in control cone versus cSCC tissue isolated from patient skin. “Cone” refers to patient-matched skin juxtaposed to tumor isolated during Mohs surgery. Each lane represents a specimen from a different patient (N = 3). kDa, kilodalton. In the dot plots below, each protein signal was normalized to its corresponding GAPDH signal. NLRP3 and ASC levels were higher in cSCC versus cone, but these differences did not reach statistical significance. However, cleaved IL-1β was significantly higher in cSCC. P-values, p > 0.05 (ns, not significant), p < 0.05 (*), were calculated using unpaired student’s *t*-tests. **D)** Representative immunofluorescence images of primary mouse ASC-citrine fibroblasts treated with vehicle (PBS), 1 µg/mL lipopolysaccharide (LPS), or 1 or 3 mM adenoside triphosphate (ATP) and stained for ASC (pink) and DNA (blue). Yellow arrows indicate ASC-citrine specks that colocalize with endogenous ASC. Scatter dot SuperPlots on the right show quantification of ASC-citrine specks (defined by specks positive for both citrine and ASC) in each treatment group. N = 4 mice per dot plot (fibroblasts were freshly isolated from separate mice). Individual values within each n are plotted in the same color. P-values, p > 0.05 (ns, not significant), p ≤ 0.001 (***), p ≤ 0.0001 (****), were calculated using ordinary one-way ANOVA with Tukey’s multiple comparison tests.

